# Zinc supplementation induced transcriptional changes in primary human retinal pigment epithelium: a single-cell RNA sequencing study to understand age-related macular degeneration

**DOI:** 10.1101/2022.09.01.504514

**Authors:** Eszter Emri, Oisin Cappa, Caoimhe Kelly, Elod Kortvely, John Paul SanGiovanni, Brian McKay, Arthur A Bergen, David A Simpson, Imre Lengyel

## Abstract

Zinc supplementation had been shown to be beneficial to slow the progression of age-related macular degeneration (AMD). However, the molecular mechanism underpinning this benefit is not well understood. In this study, we used single-cell RNA sequencing to identify transcriptomic changes induced by zinc supplementation in human primary retinal pigment epithelial (RPE) cells in culture. The RPE cells were allowed to mature for up to 19 weeks. After one or 18 weeks in culture, we supplemented the culture medium with 125 μM added zinc for one week. During maturation RPE cells developed high transepithelial electrical resistance, extensive, but variable, pigmentation and deposited sub-RPE material similar to the hallmark lesions of AMD. Unsupervised cluster analysis of the combined transcriptome of the cells isolated after two-, nine- and 19 weeks in culture, showed a significant degree of heterogeneity. Clustering based on 234 pre-selected RPE specific genes, identified from the literature, divided the cells into two distinct clusters we defined as more- and less-differentiated cells. The proportion of more differentiated cells increased with time in culture, but appreciable numbers of cells remained less differentiated even at 19 weeks. Pseudotemporal ordering identified 537 genes that could be implicated in the dynamics of RPE cell differentiation (FDR< 0.05). Zinc treatment resulted in the differential expression of 281 of these genes (FDR< 0.05). These genes were associated with several biological pathways including extracellular remodelling, retinoid metabolism and modulation of *ID1/ID3* transcriptional regulation, to name a few. Overall, zinc had a multitude of effects on the RPE transcriptome including a number of genes that are involved in pigmentation, complement regulation, mineralisation and cholesterol metabolism processes associated with AMD.

## 1. Introduction

The retinal pigment epithelium (RPE) is a highly polarized monolayer of cells lining the back of the eye, which provides critical support for the functioning of the adjacent photoreceptors. It is part of the outer blood-retina-barrier that regulates the transport of metabolites between the bloodstream and the neural retina. The RPE undergoes structural and functional transitions during maturation, which are essential to fulfil its biological functions [1]. Because of its critical function, the RPE has been directly implicated in several retinal diseases, most notably age-related macular degeneration (AMD). A hallmark feature of AMD is the accumulation of protein, lipid and mineral-rich deposits between the RPE and the choroidal microcapillary network [2]. Size and number of these sub-RPE deposits increase with disease progression [3]. Another hallmark is pigmentary changes associated with the RPE [4]. Both of these are linked to the progression to end-stage AMD [2a] manifested as geographic atrophy (GA), characterized by progressive degeneration and loss of the RPE layer, or as neovascular (NV) AMD that is characterized by abnormal leaky blood vessels that grow from the choroid into the sub-RPE space (Type 1), sub-retinal space (Type 2) or the retina (Type 3) [5], causing fluid accumulation and scarring [6]. Zinc is part of a nutritional supplement endorsed by the National Eye Institute (NEI) to slow the progression from mild|moderate to advanced AMD [7]. The biochemical pathways involved in these beneficial effects are not fully understood. Recent studies showed that human primary RPE cells in long-term culture model the hallmark features of AMD. RPE cell-based models develop as monolayers with tight junctions and high transepithelial resistance (TEER), extensive pigmentation, specific gene expression profiles and also sub-RPE deposits [8], many of which can be affected by zinc supplementation directly [8b, 8d]. This in vitro model system can be manipulated experimentally and interrogated longitudinally under conditions resembling health and disease. In this study, we identified dynamic changes in gene expression and the effects of acute (1 week) zinc supplementation using single-cell RNA sequencing (scRNA-Seq). Our results elucidate several specific pathways involved in the maturation of RPE to a stage that develops hallmark changes of AMD (sub-RPE deposition and pigmentary changes) and how these are modified by zinc supplementation.

## 2. Results

### 2.1. Maturation of RPE cells in culture

Primary human fetal RPE cells from a single donor were cultured for 2 weeks (2W=short-term), 9 weeks (9W=medium-term) and 19 weeks (19W=long-term) (Supplementary Figure 1 A). We used culture conditions that, in our hands, reproducibly recapitulated key aspects of RPE cells in previous studies [8b-d]. As time in culture increased, RPE cells developed pigmentation, hexagonal morphology (Supplementary Figure 1 B) and a progressively increasing epithelial barrier function were observed (112.9±3.9 Ohm*cm2 at 2W, 195.2±16.6 Ohm*cm2 at 9W, and 201.36 ±49 Ohm*cm2 in 19W in culture). The cell cultures also began accumulating sub-RPE deposits (Supplementary Figure 1 C). To identify the transcriptomic profiles of RPE cells at the three time points, we collected cells from 3 wells at 2W and 19W and 2 wells at 9W in culture (see Materials and Methods for detail). Approximately 3000-4000 cells were captured from each well and processed on the 10x Genomics Chromium v1.3 platform, with transcriptomes generated for a total of 30,000 cells.

### 2.2 Cluster analysis of the scRNA-Seq data identifies significant heterogeneity of RPE cells

#### Unsupervised clustering analysis

To ensure equal representation from all conditions, all samples were downsampled to include an equal number (1000) of randomly selected cells in Seurat 3.1 [9]. Based on 3,417 differentially expressed transcripts (Supplementary Table 1), the cells were automatically allocated into thirteen clusters and visualized on a Uniform Manifold Approximation and Projection (UMAP) plot (Figure 1 A).

**Figure 1.:**
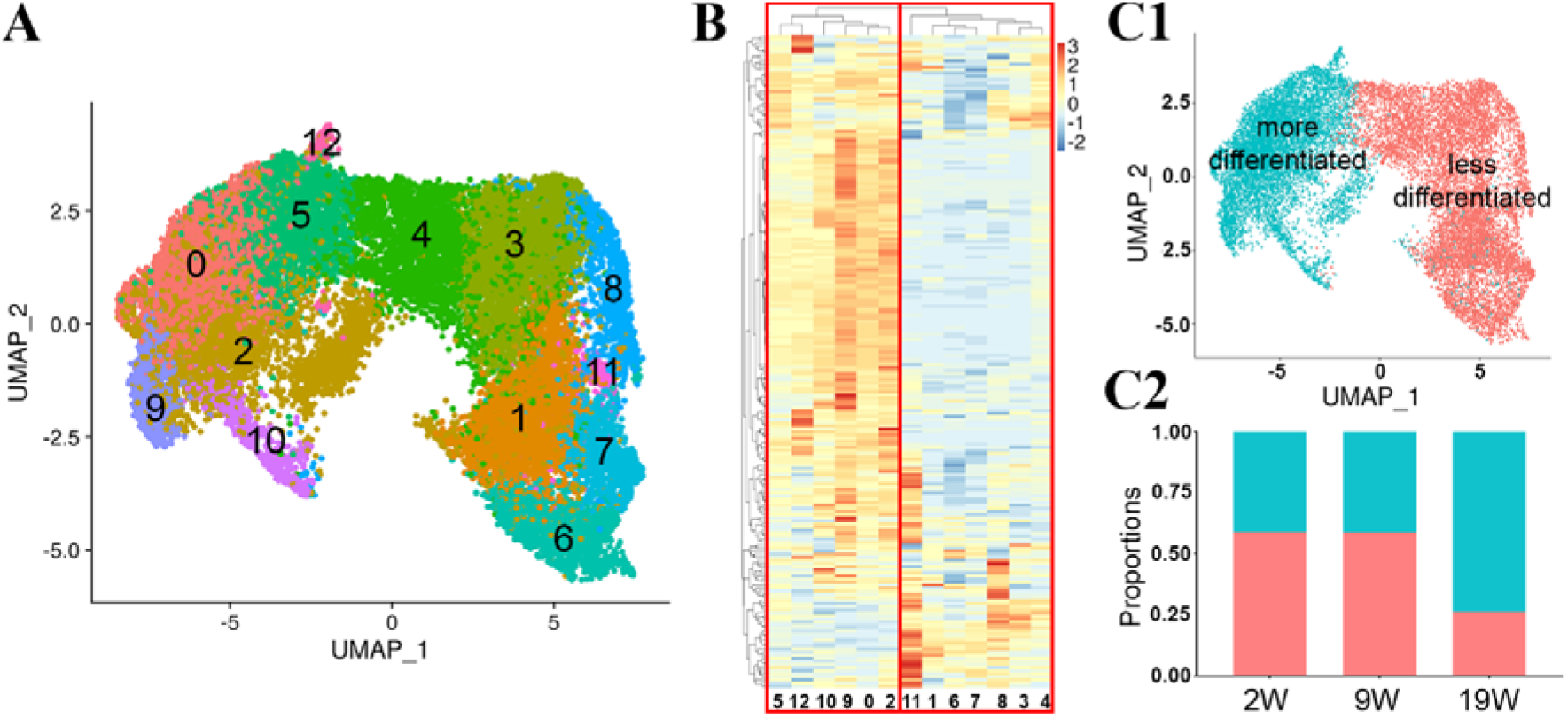
Heterogeneity of cultured human primary fetal RPE cells revealed by single-cell transcriptome analysis. (A) UMAP reduced dimensionality plot labelled with the thirteen clusters identified within the RPE cells. (B) Hierarchical clustering based on RPE-specific genes showed that the thirteen clusters partitioned into two distinct branches. (C1) UMAP replotted to indicate more (green) and less (orange) differentiated RPE cells. (C2) The proportion of more differentiated cells increased after 19 weeks in culture (blue represents more differentiated cells; orange less differentiated cells). 2W – two weeks; 9W – nine weeks; 19W – 19 weeks in culture.

The lists of cluster-specific ‘marker’ genes were input into the GeneAnalytics. The gene set analysis tool identified significant cluster-specific canonical pathways [10], labelled as superpathways for the functional analysis of the cell populations. Supplementary Table 2 contains information on the ‘marker’ genes, numbers of enriched pathways and matched number of genes to the total number of genes in a pathway for each clusters. The software assigned the 312 genes in Cluster 0 to 18 superpathways, with respiratory electron transport and heat production of uncoupling proteins, metabolism and visual cycle among the top five hits. The 205 genes in Cluster 1 were assigned to 44 superpathways, with degradation of extracellular matrix, ERK signalling and phospholipase C pathway amongst the top five hits.

The 270 genes in Cluster 2 were associated with 19 superpathways with metabolism, respiratory electron transport, heat production of uncoupling proteins, and visual cycle amongst the top five hits. In cluster 3, the 210 differentially expressed genes were associated with 76 superpathways with cytoskeletal signalling, ERK signalling and focal adhesion among the top five hits. The 87 differentially expressed genes in Cluster 4 were associated with 29 super-pathways with cytoskeletal signalling, ERK signalling and integrin signalling among the top five hits. The 30 differentially expressed genes in Cluster 5 were associated with two superpathways: melanin biosynthesis and tyrosine metabolism. The 270 differentially expressed genes in Cluster 6 were associated with 50 superpathways, degradation of extracellular matrix, metabolism of proteins, and cell adhesion and ECM remodelling amongst the top five hits. In cluster 7, 313 differentially expressed genes were associated with 110 superpathways with cytoskeletal signalling, ERK signalling and degradation of extracellular matrix among the top five hits. The 303 differentially expressed genes in Cluster 8 were associated with 61 superpathways, degradation of extracellular matrix, ERK signalling and phospholipase C pathway among the top five hits. In cluster 9, we identified 302 differentially expressed genes associated with 21 superpathways with metabolism, visual cycle and copper homeostasis among the top five hits. In cluster 10, 198 differentially expressed genes were associated with 24 superpathways with metabolism, visual cycle and oxidative stress among the top five hits. The 807 differentially expressed genes in Cluster 11 were associated with 86 superpathways, degradation of extracellular matrix, protein processing in the endoplasmic reticulum, and cytoskeletal signalling amongst the top five hits. Finally, in cluster 12, 164 differentially expressed genes were associated with 11 superpathways with organelle biogenesis and maintenance, intraflagellar transport and mitotic cell cycle among the top five hits. The identification of thirteen clusters shows that cells in culture are not homogenous.

#### Hierarchical clustering analysis using markers of mature RPE cells

We aimed to identify which of the unsupervised clusters most resemble mature RPE. We separated cells that were deemed to be more differentiated based on the expression of 213 RPE-specific genes we identified from several publications [11] (Supplementary Table 3). Hierarchical clustering based on the gene list divided the 13 clusters into two distinct groups (Figure 1 B). We annotated clusters 0, 2, 5, 9, 10 and 12 as ‘more differentiated’ RPE cells, and the remaining clusters (1, 3, 4, 6, 7, 8 and 11) ‘less differentiated’cells (Figure 1 B). We use the terms ‘more differentiated’ and ‘less differentiated’ from this point forward. The more and the less differentiated cells are clearly separated on the original UMAP (Figure 1 C1). We calculated the proportion of more and less differentiated cells at the 2W, 9W and 19W. Interestingly, nearly half of the cells were more differentiated even as early as 2W or 9W in culture (2W=41%, 9W=41%). By 19W the proportion of the more differentiated cells increased to 73% (Figure 1 C2), with 27% of the cells remaining less differentiated. Supplementary Table 4 lists the genes that define the more and less differentiated groups. The expression levels of three highly expressed representative genes from each group are presented as violin plots and UMAP plots in Supplementary Figure 2, highlighting the enrichment but not the exclusive presence of these genes in one or the other group.

Next, we tested whether the protein products of the genes that distinguish more and less differentiated cells show differential expression. One of the highly expressed mRNAs in the less differentiated cells was Collagen Type I alpha 1 chain (*COL1A1*), a fibril-forming collagen. The RPE secretes the protein encoded by this gene and it is found in the sub-RPE space [12]. We found that the expression of COL1A1 gradually increased in the less differentiated group and decreased in the more differentiated group (Figure 2 A). In contrast, Retinoid Isomerohydrolase (*RPE65*), a visual cycle component marker for differentiated RPE, was mainly expressed in the more differentiated group (Figure 2 A). Both genes were also expressed in the other group but at a low level in the opposing groups (Figure 2 B).

**Figure 2.:**
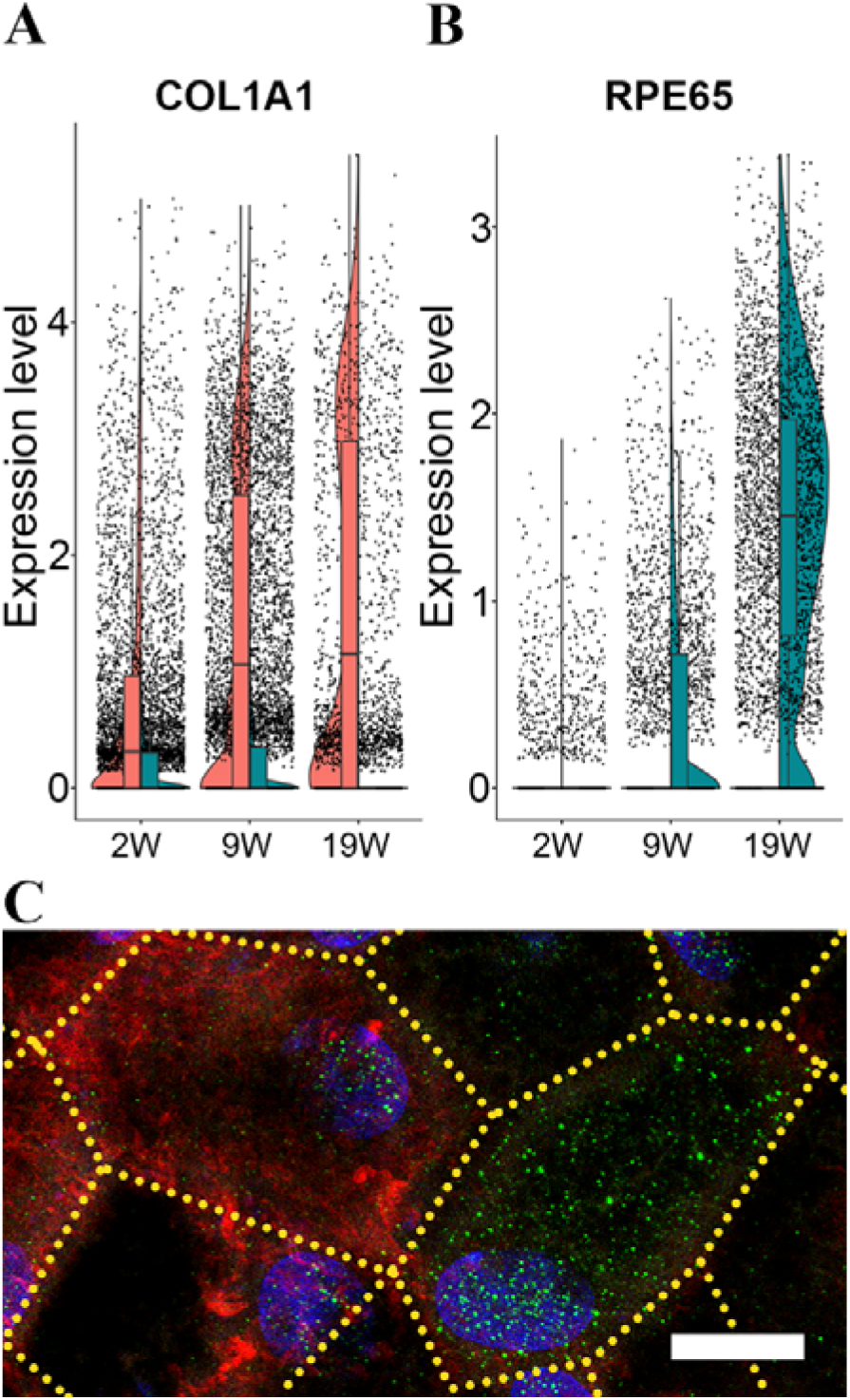
Gene expression pattern of more and less differentiated cell populations of in vitro RPE over time. Top markers of the more and less differentiated single-cell populations. Gene expressional change of COL1A1 (A) and RPE65 (B) over time at more or less differentiated cell population level, representative image of their protein expression pattern in RPE flatmounts (green: *RPE65*, red: *COL1A1*) (C). Scale bar is 10 um. 2W – two weeks; 9W – nine weeks; 19W – 19 weeks in culture.

Next, we determined the immunolocalization of the COL1A1 and RPE65 proteins in the 19W old RPE monolayer. In line with the gene expression results, the cells with a strong *RPE65* immunolabelling also had weak intracellular immunoreactivity for *COL1A1* proteins, and cells with strong immunolabelling for *COL1A1* showed weak labelling for *RPE65* (Figure 2 C; green: *RPE65*, red: *COL1A1*). Immunolabeling of *COL1A1* is also present in the sub-RPE space. This extracellular immunoreactivity gradually increased with time in culture (Supplementary Figure 3), suggesting that the secreted *COL1A1* accumulates as part of the developing extracellular sub-RPE material (Supplementary Figure 3).

As we identified more and less differentiated RPE cells in our hfRPE, we investigated whether more and less differentiated cells are also present in RPE cells directly isolated from human eyes. We used two independent previously published datasets: The scRNA-Seq data obtained from human embryos [13] or adult human eyes [14] (Supplementary Figure 4). We applied our cell grouping strategy based on the aforementioned 213 RPE specific signature genes (Supplementary Table 2). Indeed, our analysis showed that both the embryonic RPE (Supplementary Figure 4 A1 and A2) and adult RPE (Supplementary Figure 4 B1 and B2) could be classified into more and less differentiated cell populations. Of note, the number of cells analysed in the publication using adult RPE [14], was relatively low, hence clusters were less well separated.

### 2.3. Pseudotemporal ordering of the expressed RPE genes

To identify the genes associated with transitioning from the less to the more differentiated cells, we performed a pseudotemporal ordering of our scRNA-Seq transcriptome profile using Monocle3 (Figure 3). This unsupervised analysis identified a main trajectory with 11 nodes (Figure 3 A1). Based on the original cluster analysis depicted in Figure 1, node 1 corresponded to the less and node 10 to more differentiated cells (Figure 3 A2). The main trajectory was correlated with 537 variably expressed genes. Based on their pseudotemporal expression profile, these clustered into seven modules (Figure 3 B1; Supplementary Table 5). Modules 2 and 5 contained 175 genes with high expression at the early stages of the trajectory that gradually declined towards the end of the trajectory. GeneAnalytics identified 62 potential significant superpathways associated with these genes (Supplementary Table 6). Degradation of extracellular matrix, focal adhesion, and cell adhesion-endothelial cell contacts was amongst the top five ranked pathways. Modules 3, 4 and 6 contained 172 genes. These gradually increased towards the late stages of the trajectory. GeneAnalytics identified nine potential superpathways defined by these genes, including transport of glucose, metabolism and visual cycle among the top five hits (Supplementary Table 6). Modules 1 and 7 contained 190 genes. The expression of these genes transiently increased to a maximum at the middle of the trajectory. These identified sixteen superpathways with degradation of extracellular matrix, ERK signalling and cytoskeleton remodelling among the top five hits (Supplementary Table 6).

**Figure 3.:**
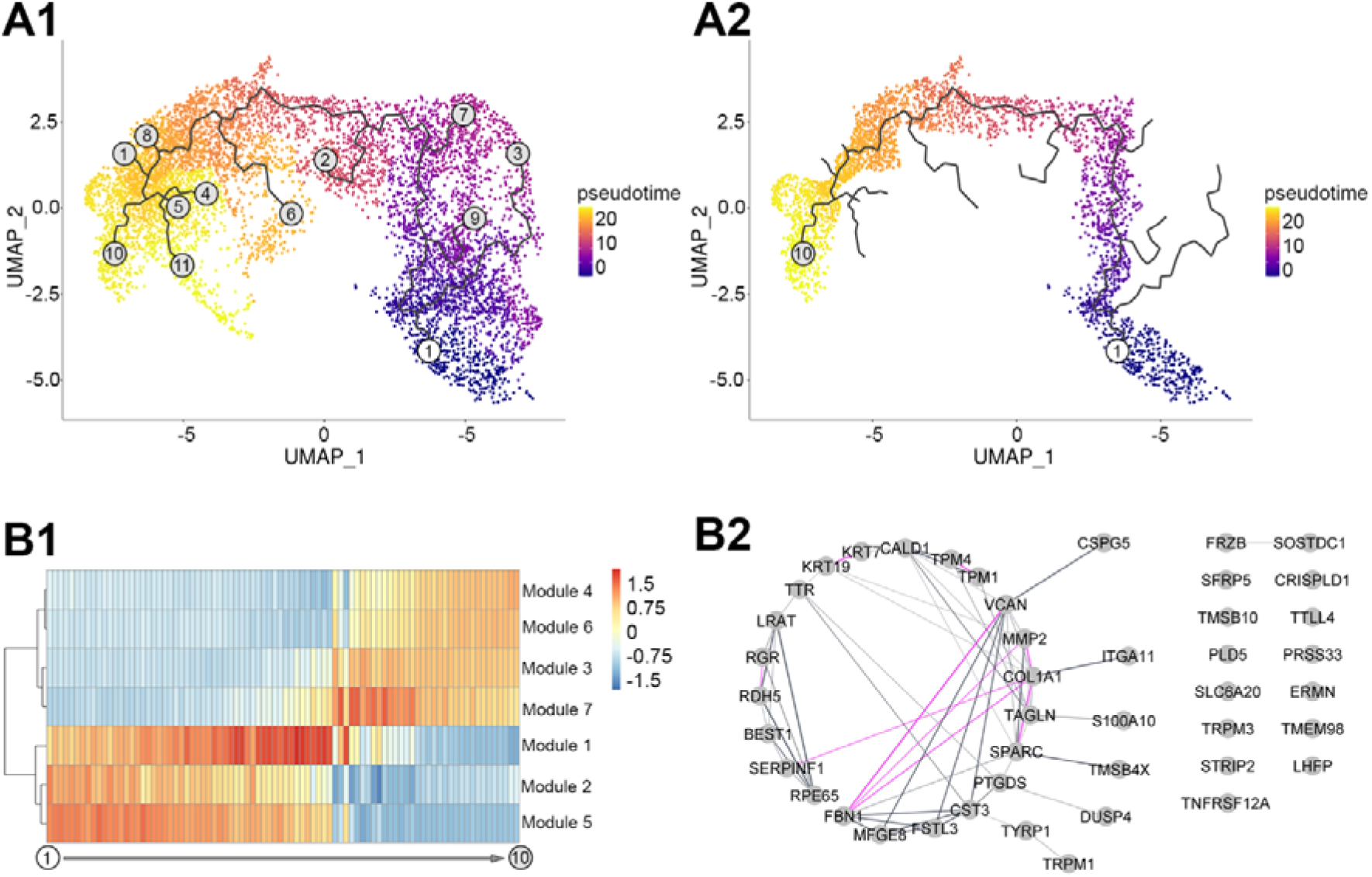
Dynamic changes of RPE over time in vitro. Identified trajectories (A1) and the highlighted main trajectory (A2) of in vitro RPE transcriptome over pseudotime (A), heatmap visualisation of the expression patterns the genes correlated with the main trajectory grouped into modules (B1) and network representation of the genes showing highest correlation with main trajectory of RPE culture pseudotime above the cut-off value of Moran I using Gene Analytics (B2, grey lines represent validated connection via text-mining, database information and co-expression, pink lines represent experimentally validated network connection, the thickness of lines indicates the strength of data support).

To identify potential transcriptional regulators of the pseudotemporal trajectory, GO term analysis was carried out in GeneAnalytics. This identified transcriptional regulator activity in two genes, *ID1* and *ID3*, belonging to the combined module 1 and 7, representing the transitional phase on the pseudotemporal trajectory.

Next, we examined the potential relationships between the most highly significantly correlated in the main trajectory using a cut-off value of Moran I = 0.5 (see Material and methods section). We identify 44 genes strongly influencing the main trajectory. Using GeneAnalytics in combination with STRING database and Cytoscape we found that these genes do not appear to be randomly distributed. Thirty-one out of 44 genes showed a significant biological connection, validated via text-mining, database information, co-expression, or experimental evidence. (*p*<1.0e-16; Figure 3 B2). The 44 genes were associated with 6 potential superpathways, including visual cycle, degradation of the extracellular matrix and cell adhesion-extracellular matrix remodelling as the top hits (Supplementary Table 6).

### 2.4. Acute zinc supplementation has a multitude of effects on transcription in RPE cells

#### Transcriptional changes in response to Zinc supplementation

To determine whether the beneficial effects of zinc described by previous studies [7, 8b, 8d, 15] are mediated, at least in part, through changing transcription we treated our RPE cultures for one week with a zinc-supplemented medium, as described earlier [8b]. This acute zinc supplementation was carried out on less differentiated cells starting at the end of the first week in culture, and then cells were harvested at the end of the 2W or more differentiated cells at the end of the 18^th^ week and cells were harvested at the end of the 19W. Gene expression changes with zinc supplementation were compared to cells in culture without zinc supplementation for either 2W or 19W. Cells with and without zinc supplementation were clustered using the process used in Figure 1 C1. While acute zinc supplementation did not noticeably change the proportion of the more and the less differentiated cells (Figure 4 A), it significantly changed the expression of 472 genes in the more differentiated cells (Figure 4 B1) and 149 genes in the less differentiated cells (Figure 4 B2) at the two-week time point (Supplementary Table 7).

**Figure 4.:**
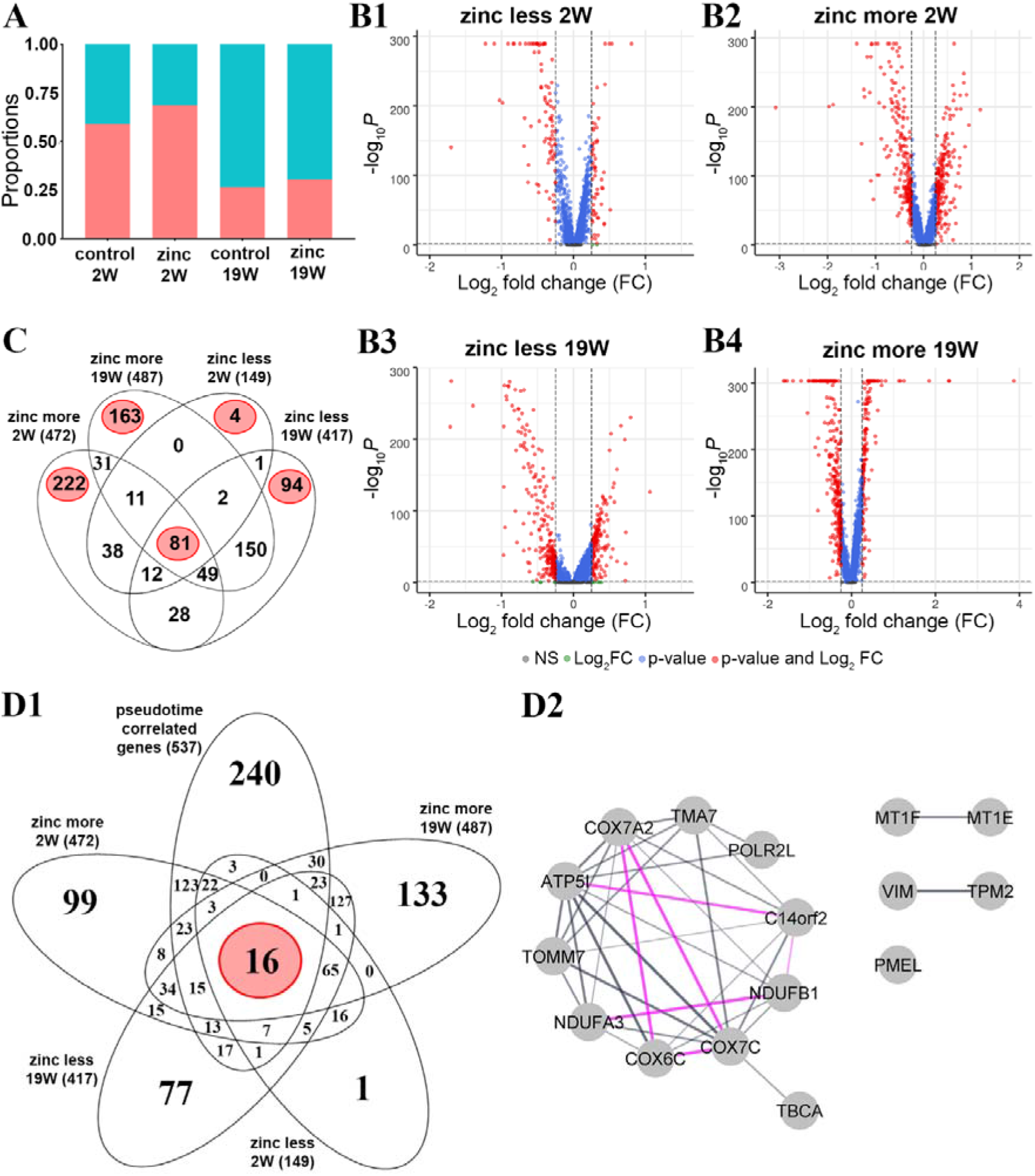
Impact of acute zinc supplementation on in vitro RPE at single-cell level. Relative proportion of less and more differentiated cells over time following acute zinc treatment (A), volcano plot visualisation of the differentially expressed genes in less (B1) and more (B2) differentiated cells in two week-culture and in less (B3) and more (B4) differentiated cells in 19 week-culture, number of differentially expressed genes overlapped amongst the different cell types and culture times (C), overlap between pseudo-time correlated genes and differentially expressed genes following acute zinc supplementation (D1) and a network representation of the 16 overlapping genes using Gene Analytics in combination with String and Cytoscape (D2), in which grey lines represent validated connections via text-mining, database information and co-expression, pink lines represent experimentally validated network connection and the thickness of lines indicates the strength of supporting data.

At 19W, zinc altered the gene expression of 487 genes in the more differentiated cells (Figure 4 B3) and 417 genes in the less differentiated cells (Figure 4 B4) (Supplementary Table 7) (logFC> 0.25, adjusted p-value< 0.05). We displayed the four datasets in a 4-way Venn-diagram to further analyze specific temporal zinc-induced gene expression changes (Figure 4 C). We found 81 overlapping genes differentially expressed under all four conditions. Two-thirds of these 81 genes were identified as housekeeping genes by GeneAnalytics confirming previous studies showing that zinc plays a role in regulating cellular homeostatic processes [16]. Relevant proteins include metallothioneins (MT1E, MT1F and MT1X) that act as essential stress proteins to regulate immune homeostasis. In the more differentiated cells, 222 uniquely affected genes were at 2W and 163 in the 19W (Figure 4 C). In the less differentiated cells, only four genes were specifically affected by zinc supplementation at 2W and 94 genes at 19W (Figure 4 C).

At 2W, we identified superpathways only in the more differentiated cells; these were cytoskeleton remodelling, focal adhesion and degradation of extracellular matrix among the top five superpathways (Supplementary Table 8). At 19W, in the less differentiated cells, we identified presenilin signalling, SMAD signalling and antigen presenting-cross presentation amongst the top five superpathways (Supplementary Table 8), while in the more differentiated cells, we identified metabolism, ferroptosis and protein processing in the endoplasmic reticulum amongst the top five superpathways (Supplementary Table 8). Information on the magnitude and direction of zinc-associated change in transcript abundance of these gene lists are in Supplementary Table 7, and the analysis of these five gene lists by GeneAnalytics to identify superpathways are listed in Supplementary Table 8.

#### Influence of Zinc on Transcription Dynamics

We next determined the overlap between the 537 genes identified in the main trajectory in the pseudo-temporal analysis (Figure 3 A2; Supplementary Table 5) and the list of the differentially expressed genes following the acute zinc supplementation (Figure 4 D1; Supplementary Table 7). This comparison identified 16 common genes (Supplementary Table 9). Using GeneAnalytics in combination with STRING database and Cytoscape we found that these 16 genes show significantly (*P*-value < 1.0e-16) more interactions than expected, validated by text-mining, database information, co-expression, and experimental evidence (Figure 4 D2) that relates to the respiratory electron transport and response to metal ions as biological function (Supplementary Table 9).

### 2.5. Sub-RPE deposition-related gene expression pattern depends on maturation state and zinc supplementation

Our hfRPE culture developed sub-RPE deposits even without photoreceptors and the supporting choriocapillaris (Supplementary Figure 1C). This allowed us to analyze the expression of genes potentially involved in the sub-RPE deposit formation process. We compiled lists of genes associated with various aspects of sub-RPE deposit formation and analyzed the changes in expression throughout cell maturation and zinc supplementation (Supplementary Table 11). Some genes belong to more than one gene list (Figure 5).

**Figure 5.:**
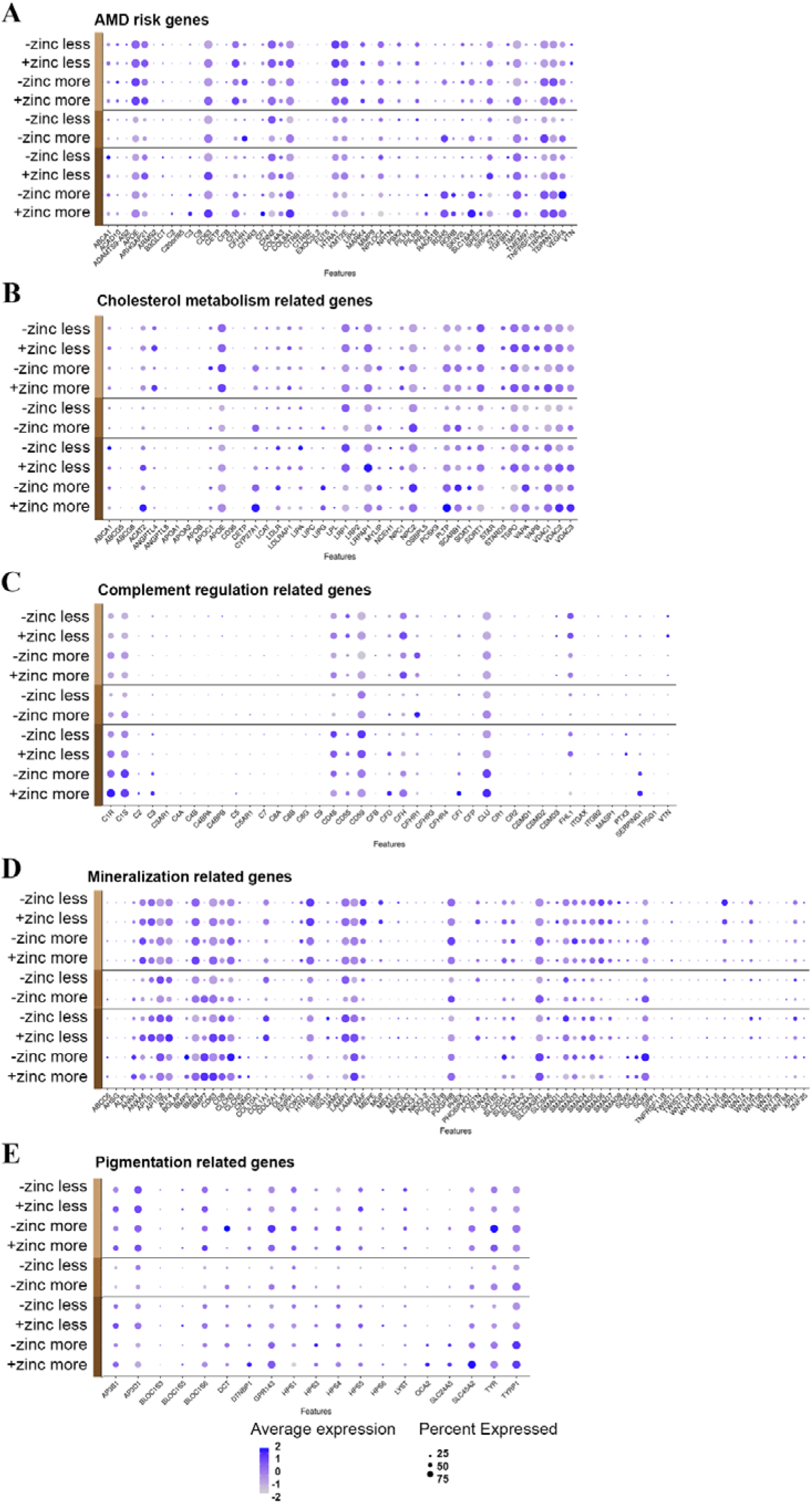
Distribution of age related gene expression in RPE over time in vitro. AMD risk genes (A), cholesterol metabolism-related genes (B), complement regulation-related genes (C), mineralisation-related genes (D) and pigmentation-related genes (E). Light brown represents two weeks, medium brown represents nine weeks, dark brown represents 19 weeks in culture.

#### Genelist 01

contains 55 genes previously genetically associated with AMD (Figure 5 A; Supplementary Table 11 from [17]. We found that 52 out of the 55 genes in AMD risk-associated risk loci were expressed in our RPE model (Supplementary Table 11). Some genes were expressed higher at 2W, like *CFHR3, LIPC, SYN3* and *VTN*, while others were expressed higher at 19W, like *ARHGAP21, RDH5, SKIV2L, SRPK2, TGFBR1* and *TRPM3*. Amongst the genes that were expressed higher in less differentiated cells were *CFHR3, LIPC, TGFBR1* and *VTN*. In more differentiated cells, we found higher expression of *PRLR, RDH5, RORB, SLC16A8, SPEF2* and *VEGFA*. From these 52 genes, *CFH, COL8A1, CD63, TSPAN10, APOE, TIMP3* and *SLC16A8* were significantly upregulated, while *CFHR1, VEGFA, TRPM3* and *RDH5* were significantly down-regulated in response to acute zinc supplementation (Supplementary Table 7).

#### Genelist 02

contains 66 complement-regulation-related genes. Several complement proteins have been implicated in AMD and are found in sub-RPE deposits (Figure 5 B, Supplementary Table 11 [18]). 41 out of the 66 identified complement genes were expressed in our hfRPE cultures, most showing low expression levels (Supplementary Table 11). The genes that were expressed higher in 2W cultures were *C4B, C4BPB, C8B, C8G, CFHR3, CFP, CSMD1, CSMD3, TPSG1* and *VTN*, while the genes expressed higher in 19W cultures were *C1S, CD55, CD59* and *PTX3*. Among the genes expressed higher in less differentiated cells were *C4BPA, C4BPB, ‘C8G, CFHR3, CFP, CR2, CSMD1, FHL-1, ITGB2, PTX3* and *VTN*. The expressions of C2, C4A, C5, CR1 and SERPING1 were higher in the more differentiated cells. *CFH* and *C1R* were significantly upregulated, while *CFHR1* and *CLU* were significantly down-regulated in response to Zinc supplementation (Supplementary Table 7).

#### Genelist 03

contains cholesterol metabolism-related genes (Figure 5 C, Supplementary Table 11 [19]). 42 out of the 51 identified genes were expressed in hfRPE (Supplementary Table 11). The genes that were expressed higher at 2W were *ABCG5, ANGPTL8, APOA1, CD36, LIPC, and STAR*, while the genes expressed higher at 19W were *APOC1, LDLRAP1* and *NPC1*. Among the genes expressed higher in less differentiated cells were *CD36, LIPC, PCSK9* and *STAR*. In the more differentiated cells, the expression of *CYP27A1, LIPG, LPL* and *PLTP* were higher. *PLTP, ANGPTL4, APOE, LRPAP1, VDAC2* and *TSPO* were significantly upregulated, and *NPC2* was significantly down-regulated in response to zinc supplementation. Interestingly, *CYP27A1* showed up-regulation significantly at 2W and significant down-regulation at 19W in response to zinc supplementation (Supplementary Table 7).

#### Genelist 04

contains mineralization-related genes (Figure 5 D, Supplementary Table 11 [20]) that could be associated with the inorganic hydroxyapatite component of sub-RPE deposits. Eighty out of the identified 99 calcification-related genes were expressed in the RPE (Supplementary Table 11). The genes that were expressed higher at 2W were *COL10A1, NKX3-2, PHEX, SPP1, TNFRSF11B*, and *WNT7B*, while others were expressed higher at 19W, including *AP1S2, BMP2, CLCN3, LAMP1, POSTN, SMAD1* and *SOX9*. Among the genes expressed higher in less differentiated cells were *AP1S1, COL10A1, COL1A1, DLX5, IBSP, JAM2, LAMP1, MGP, MYORG, NKX3-2, PDGFB, PHEX, POSTN and RUNX2*. The expression of ABCC6, BMP2, BMP7, CNMD, SOX6 and WNT6 were higher in the more differentiated cells. *COL1A1, POSTN, CD63, LAMP1* and *BMP4* were significantly upregulated and *SLC20A1, SOX9* and *BMP7* were significantly down-regulated in response to zinc supplementation (Supplementary Table 7).

#### Genelist 05

contains genes that are related to pigmentation (Figure 5 E, Supplementary Table 11 [4a, 21]. Pigmentary abnormalities show strong correlation with sub-RPE deposit formation and the development of AMD, and we found that 19 out of the identified 21 genes were expressed in hpRPE (Supplementary Table 11). At 2W, we found no differentially expressed genes. At 19W, however, we found that *AP3B1, AP3D1, BLOC1S6, HPS5, HPS6*, and *SLC24A5* were expressed higher. There were no highly expressed genes in less differentiated cells. In the more differentiated cells, the expression of *OCA2* and *SLC24A5* was higher. *TYR, TYRP1* and *DCT* were significantly downregulated in response to Zinc supplementation in our acute treatment (Supplementary Table 7).

## 3. Discussion

The RPE plays a pivotal role in maintaining the health of the retina, and changes in RPE function have been linked to the development and progression of AMD [22]. Optimal zinc balance is key for RPE function [23], and deficiency in zinc contributes to AMD pathogenesis [24]. Based on these findings, it has been suggested that zinc supplementation can slow the progression of AMD [24–25], although the mechanism of this beneficial effect is not fully understood [26]. In this study, we used primary human fetal RPE cells and scRNA-Seq analysis to identify the transcriptomic changes and biologically plausible molecular pathways involved in the maturation of the RPE and the changes associated with zinc supplementation. The specific transcriptional changes and molecular pathways identified provide an improved understanding of RPE cell maturation and insight into how the function of RPE might be affected by acute zinc supplementation which has relevance for the progression of AMD.

### 3.1. Study rationale

Maturation of RPE cells is key to developing appropriate morphology, pigmentation [27] and the production of key signature proteins that determine the function of these cells [28]. Different studies use a variety of sources to study RPE maturation and function, ranging from the immortalized ARPE-19 cells [29] to induced pluripotent stem cell-derived RPE [30] and primary porcine [8c] or human RPE [31]. As with all model systems, cellular models for RPE must replicate the *in vivo* situation as close as possible. Recently we have shown that primary human fetal RPE cells develop the most critical features of native RPE, including the formation of pigmentation, tight junctions with high TEER values, and the expression of RPE signature genes and proteins [8b, 8c]. Most importantly, the cells in culture can lay down sub-RPE deposits, a hallmark feature of AMD [8b, 8c]. Despite demonstrating these in vivo-like features, the molecular signature for RPE maturation has not yet been fully explored. Previous studies have reported a variety of approaches to map molecular maturation. Earlier studies used microarrays [11c, 32] or bulk RNA sequencing. Most recently, a powerful tool capable of sequencing individual cells has been introduced. Single-cell RNA sequencing provides an unparalleled opportunity to identify cell heterogeneity [33]. Lidgerwood et al. [34] used pluripotent stem cell-derived RPE to analyze transcriptomic changes after 1 month or 12 months in culture and analyzed these separately, then combined the data. In a subsequent study, the same group combined scRNA-Seq and proteomics in iPSC cells obtained from individuals with or without AMD to identify regulations in geographic atrophy [35]. Exciting opportunities are presented by scRNA-Seq studies using freshly isolated RPE from human eyes. RPE cells from both fetal and adult human eyes were analyzed in previous studies [13] and [14], respectively. Although both studies used a limited number of cells, they provide an invaluable addition to cell culture-based observations. In our study, we used primary fetal RPE cells that recapitulated features of RPE cells in vivo (Supplementary Figure 1). Despite their fetal origin, these cells developed sub-RPE deposits and varied pigmentation, suggesting that they recapitulate the hallmarks of AMD (Supplementary Figure 1) despite the relatively short time in culture (19W).

### 3.2. Heterogeneity of RPE cells

The generation of scRNA-Seq data from a large number of cells allowed us to confidently determine that there is a significant degree of heterogeneity between the cells. A key observation was that some RPE cells could develop into more differentiated cells even after two weeks in culture, but even after 19 weeks, we still observed less differentiated cells (Supplementary Figure 1). Heterogeneity of RPE had been reported after multiple passages and over years in culture, [36] (Supplementary Figure 1), reflecting what had been reported for RPE *in vivo* [37] and *in situ* [36]. Despite the long-lasting heterogeneity, the melanosome precursor *PMEL17* was expressed in both less and more differentiated cells. In fact, from the 19 pigmentation-related genes expressed in our cells, the only transcript that showed elevated expression in the more differentiated RPE cells were *OCA2* and *SLC24A2* (Supplementary Figure 2 B1, Figure 5 E, Supplementary Table 11), suggesting that all cells could become pigmented [21a, 38].

*COL1A1* was amongst the top transcripts in the less differentiated cells, and immunoreactivity of COL1A1 protein was able to distinguish the less differentiated cells from the more differentiated cells that express *RPE65* gene highly and are immunopositive for the RPE65 protein (Figure 2 C). Immunoreactivity to the COL1A1 protein gradually increased in the sub-RPE space with time in culture (Supplementary Figure 3), suggesting that the half-life of this extracellular matrix protein is long in our culture system. This increase in sub-RPE COL1A1 may correspond to the role this protein plays in forming the extracellular matrix of Bruch’s membrane [39]. Other collagens were also expressed highly in the less differentiated cell population (Supplementary Table 4), reflecting their reported involvement in increased attachment, and spread of RPE cells [40]. The only highly expressed transcript for collagen in the more differentiated cells was *COL8A1* (Supplementary Table 4). The COL8A1 protein is a component of basement membranes in the eye and contributes to the formation of the basement membrane of RPE [11a, 41] and a genetic risk variant of AMD [42]. The findings on COL1A1 and RPE65 might be mechanistically important: the mature RPE cells (*RPE65* expressing) could enable performance of the visual cycle, while the less differentiated cells (*COL1A1* expressing) can support the formation of ECM throughout life.

### 3.3. Transition from less to more differentiated RPE

As the more and the less differentiated cells are present at all the three-time points, we combined the scRNA-Seq data from the 3 time points and analyzed these datasets together, an approach different from a previous study [34]. This integrated approach helped us to identify a pseudo-temporal trajectory of gene expression from less- to more-differentiated cells (Figure 3 A1). This approach identified a well-defined main trajectory (Figure 3 A2). The top genes with the highest score in the main trajectory were associated with regulating the visual cycle (*RPE65, LRAT, TTR, RDH5*) (Supplementary Table 6). Transcriptomic analysis of the bulk RNA isolated from RPE cells from ageing human donor eyes recently reported a positive feedback mechanism between up-regulation of visual cycled genes and the accumulation of retinoid by-products [43]. As visual cycle-related bisretinoids are constituents of the accumulating lipofuscin in RPE [44], this up-regulation could eventually lead to AMD-like pathogenesis [45] in this cell culture model. Indeed, there are ongoing clinical trials for visual cycle modulators as therapeutic options for AMD [46], and our cell culture model has the potential to serve as a preclinical tool for testing novel compounds.

### 3.4. Genes involved in transitioning RPE from less to more differentiated cells

The genes associated with the main trajectory could be clustered into seven modules based on their transcriptional change along the pseudotemporal trajectory (Supplementary Table 5). The transcripts whose expression is transiently up-regulated on the pseudotemporal trajectory likely represent the genes mediating the transition from the less to the more differentiated cells (Supplementary Table 5). These genes were associated with cellular and extracellular remodelling and metabolic pathways (Supplementary Table 6). Therefore, our data support the hypothesis that extracellular matrix remodelling of the Bruch’s membrane could become a therapeutic target to combat RPE loss [47] due to topographic changes in the RPE-Burch’s membrane interface [40]. Alterations of the extracellular matrix may impact immune response as well as the secretion of pro-inflammatory cytokines, such as MCP-1, and IL-8 [40] and promote sub-RPE deposit formation [48]. Our data highlights potential molecular targets to achieve a regulation of this process.

Among the transiently expressed genes, we identified *ID1* and *ID3* (Supplementary Table 6). The corresponding helix-loop-helix (HLH) proteins form heterodimers with members of the basic HLH family of transcription factors, inhibiting DNA binding and preventing the formation of active transcriptional complexes [49]. ID proteins promote cell cycle progression and cell migration and restrict cellular senescence and the differentiation of a number of progenitor cell types [50]. Recent results indicate that the expression of ID family proteins may play an important role in regulating retinal progenitor cell proliferation and differentiation [51]. ID genes and proteins showed increased expression levels in the retina at embryonic and early postnatal stages and declined in the adult [51]. ID protein expression is silenced in many adult tissues but is re-activated in diverse disease processes [50b, 52]. ID proteins appear to play a crucial role in the angiogenic processes and it was proposed that inhibition of expression and/or function of ID1 and ID3 may potentially be of therapeutic value for conditions associated with pathological angiogenesis [53]. In fact, deletion of Id1/Id3 reduced ocular neovascularization in a mouse model of neovascular AMD [49]. In conclusion, drugs targeting ID1/ID3 could modulate RPE maturation and pathological changes in AMD.

### 3.5. Response to acute zinc supplementation

Treatment with zinc has been reported to prevent progression to advanced AMD (for review see: [54]) at least partly due to a direct effect of zinc on the RPE [8b, 55]. In previous *in vitro* studies, we investigated long-term supplementation with zinc and found altered selective gene expression and protein secretion as well as increased pigmentation and barrier function [8b, 8d]. We identified several molecular pathways, such as cell adhesion/polarity, extracellular matrix organization, protein processing/transport, and oxidative stress response, involved in the beneficial effects of chronic zinc supplementation to the RPE. However, these studies were unable to address the complexity associated with cell heterogeneity and detailed temporal changes. We were particularly interested in exploring how zinc supplementation could affect the less and more differentiated cells short-term to understand the potential to develop a more targeted intervention through supplementation.

To decipher the effects of acute zinc supplementation RPE cells were treated with elevated zinc for 1 week following the protocols we used previously [8d]. We found that acute zinc supplementation induced significant changes in gene expression in both short and long-term cultures (Figure 4 B1-4) regardless of the temporal stage of the cells. We also identified 81 zinc-responsive transcripts (Figure 4 C) that were common amongst all groups. These transcripts were enriched in housekeeping genes and contained transcripts for metallothioneins, ribosomal protein, and ATP synthases (Supplementary Table 8), indicating that zinc effects the cellular homeostasis of the RPE, similar to that of other systems [56]. Apart from the shared genes, there were specific changes associated with the more or the less differentiated cell groups at both 2W and 19W in culture (Figure 4 C). The four specific genes affected by short-term zinc supplementation in the less differentiated cell group (Supplementary Table 8) are genes linked to the integrity of Bruch’s membrane (COL8A1) [42], epithelial-mesenchymal transition (KRT17) [57], phagocytic activity and the rescue of the RPE (MFGE8) [58] and activity of heparan sulfate (SULF1) [11b], suggesting that zinc might influence interaction with local extracellular environment. In the more differentiated cell group in the 2W cultures, zinc affected biological processes including extracellular matrix organization, cellular polarity and visual processes (Supplementary Table 8) that are critical for supporting the photoreceptors [59].

At 19W in culture zinc affected the less differentiated cells via modulating proteolysis, DNA replication and RNA transcription and amino acid metabolisms (Supplementary Table 8) probably to mitigate oxidative stress, one of the AMD associated biological functions [60]. In the more differentiated cells in 19W in culture zinc supplementation affected several metabolic pathways (Supplementary Table 8). Dysregulation of metabolic pathways is an important contributor to AMD pathophysiology [61] and this may directly explain the benefit observed with zinc supplemented in patients in the AREDS study [7, 24, 62]. Therefore, zinc supplementation has a multitude of effects on RPE, with some specific effects depending on cell differentiation and maturity. The identification of the specific molecular changes may help redefine treatment strategies based on zinc supplementation or nutritional interventions.

### 3.6. The effects of zinc on the genes in the pseudotemporal trajectory

Earlier we identified 537 genes (Supplementary Table 5) in the main pseudotemporal trajectory (Figure 3A2). Zinc supplementation had no effect on 240 genes (Figure 4 D1). Of the remaining 297 genes 16 were housekeeping genes (Supplementary Table 8; Figure 4 C) associated with the mitochondrion, the activation of cytochrome-c oxidase and ubiquinone and response to metal ions (Supplementary table 9). This is in line with a previous observation that zinc supplementation can protect the RPE from oxidative stress-induced cell death by improving mitochondrial function [55a] and this could be behind the positive effect of zinc supplementation in the AREDS studies [7, 24, 62] or increased zinc intake through diet [15b, 63]. Metallothioneins (MT1F and MT1E) that belong to this group (Figure 4 D2, Supplementary Table 7 and 9) are well recognized mediators of zinc supplementation in the RPE [64] via mediating oxidative stress induced RPE damage [55b] and differentiation of RPE [34].

The remaining 281 genes in the main trajectory (Figure 4 D1) were associated with a variety of biological processes including extracellular matrix organization, angiogenesis, collagen fibril organization and visual perception (Supplementary Table 10). The composition of extracellular matrix has a profound effect on how the RPE attach to the Bruch’s membrane [65] and therefore modification of gene expression by zinc could directly affect sub-RPE deposit formation [48a].

We also found that acute zinc supplementation up-regulated the transcriptional regulators *ID1* and *ID3* expression, a finding that had not been reported before. In addition, in a previous study we have identified *TGFB1* as a potential upstream regulator effect of chronic zinc supplementation [8d] and we found here that *TGFB1* expression was also up-regulated by acute zinc supplementation in our current study. Therefore, we carried out an Upstream Analysis in Ingenuity Pathway Analysis (QIAGEN, Redwood City) for the 190 transiently expressed genes in the combined pseusotime correlated groups 1 and 7 (Supplementary Table 5). We identified a strong relationship for *TGFB1* (p<6.98e-19) and also for *ID1* (p<3.59e-05) and *ID3* (p<1.23e-03) as potential upstream regulators for a group of genes among the transiently expressed group. In fact, TGFB1 was an upstream regulatory element for ID1 and ID3 (Supplementary Table 12). A direct molecular link between ID1 and TGFB1 had already been suggested [66]. Therefore, the positive effects of zinc supplementation could be directly through TGFB1 signalling that involves ID1 and ID3. As the receptor of TGFB1, TGFBR1, is an AMD genetic risk variant [17] suggesting that these findings have direct relevance to further studies on AMD.

### 3.7. AMD specific gene expression changes

Based on literature searches we generated gene lists that have been shown to contribute to the pathological changes associated with AMD and we examined the effects of cell maturation and zinc supplementation on these genes (Figure 5, Supplementary Table 11). Specific attention was paid to the activation complement system and lipid metabolism related genes as these were the genetically most significantly associated pathways with AMD [17, 67]. We also scrutinized genes associated with pigmentary changes and mineralization associated genes due to their potential link with RPE function and/or sub-RPE deposit formation in AMD [2a, 20d].

Not all genes involved in complement regulation were expressed in RPE cells (Figure 5 B, Supplementary Table 11). This is perhaps not surprising, as the local activity of the complement cascade is influenced by a complicated mix of local and systemic regulatory factors, which is altered in AMD retina [1d, 68]. However, some complement genes that were expressed in the RPE were affected by acute zinc supplementation including CFH*, C1R, CFHR1* and *CLU* (Figure 5 B, Supplementary Table 7). These transcriptomic changes support our previous reports that zinc supplementation has a functional effect on CFH secretion [8d] as well as oligomerization and activity [69] and zinc levels can regulate interferon gamma systematically, that, in turn, regulates expression of complement genes [70] [71]. Apart from CFH several complement proteins can bind zinc and this binding altered their activity [72]. In addition, a network analysis has highlighted elements of the complement regulation as potential targets for nutrient-affected pathways [73]. Finally, there is also clinical evidence that zinc supplementation can indeed directly inhibit complement activation in AMD patients [68a] suggesting that modulation of the complement system could be one of the ways that zinc supplementation affects the progression to AMD.

Of the 42 genes expressed in our RPE culture associated with cholesterol metabolism (Figure 5 C, Supplementary Table 11), *ANGPTL4, LRPAP1, VDAC2, APOE, PLTP,, NPC2, TSPO* and *CYP27A1* were altered in response to acute zinc supplementation (Figure 5 C, Supplementary Table 7) [74]. These findings corroborate our previously reported effect of long-term zinc supplementation on lipid metabolism [8d]. ANGPTL4 is a lipid-inducible feedback regulator of LPL-mediated lipid uptake. However it is also a multifunctional cytokine, regulating vascular permeability, angiogenesis, and inflammation [75]. The systemic level of ANGPTL4 is associated with NV AMD [76]. Reportedly, this protein indirectly induces RPE barrier breakdown [77]. LRPAP1 is a chaperon protein, that, in general, controls the folding and ligand-receptor interactions expression of the LRP receptors [78]. Its role in RPE and AMD remains elusive. VDAC2 is a ceramide sensor integrated into the mitochondrial membrane and its function relates to regulation of mitochondrial apoptosis [79]. Increased ceramide levels affect non-polarized RPE cells found in late stages of AMD [80]. APOE, a lipophilic glycoprotein with major role in lipid transport, is one of the many constituents of the sub-RPE deposits and has been associated with increased AMD risk [81]. PLTP is a phospholipid transfer protein and is one of the main players of lipid homeostasis in ApoB-containing particles and high-density lipoprotein metabolism. PLTP plasma levels are associated with AMD [82], but their potential role in drusen formation remains elusive. NPC2 is a cholesterol transporter, effluxing cholesterol out of late endosomes in RPE. The lack of this protein is associated with age-related maculopathies [83]. TSPO is a translocator protein, transferring cholesterol from the mitochondrial outer membrane to the mitochondrial inner membrane and also plays role in oxidative stress and inflammation. It was recently implicated as a highly relevant drug target for immunomodulatory and antioxidant therapies of AMD [84]. CYP27A1 is involved in the elimination of 7-ketocholesterol from RPE, a toxic product of cholesterol autooxidation, which accumulates in drusen [85]. In summary, the aforementioned affected gene expression in response to zinc suggest, that zinc has an impact on sub-RPE cholesterol accumulation, oxidative stress, inflammation and angiogenesis via the regulation of lipid-membrane interaction, lipid transport and the elimination of toxic lipid byproducts.

In our cultures, we found 80 RPE-expressed genes associated with mineralization (Figure 5 D, Supplementary Table 11). Out of these, we found that *COL1A1, POSTN, CD63, LAMP1, BMP4, SLC20A1, SOX9* and *BMP7* were altered in response to acute zinc supplementation (Figure 5 D, Supplementary Table 7). The *POSTN* gene encodes a secreted extracellular matrix protein that functions in tissue development and regeneration and a potential anti-fibrotic therapeutic target for NV AMD [86]. CD63 is involved in the regulation of cell development, activation, growth and motility [87] and together with LAMP1, it plays a role in autophagy, exosome secretion and drusen formation [88]. BMP4 has been implicated in the disruption of RPE cell migration and barrier disruption in NV AMD [89]. The protein encoded by SLC20A1 is a sodium-phosphate symporter, which is involved in vascular calcification but not reported in association with RPE function or AMD [90]. SOX9 plays a key role in the regulation of visual cycle gene expression in RPE [91] but also plays a role in prevention against calcification [92]. BMP7 is hypothesized to be critical for differentiation of the retinal pigmented epithelium during development [93]. It also has been implicated in prevention of vascular calcification [94]. Zinc supplementation is reported to inhibit phosphate induced vascular calcification [95], but, as our results indicate, it may also have a (indirect) role in the prevention of drusen calcification.

In our cultures, the majority of pigmentation-related genes were detected and their expression level either remained constant or increased throughout the culture time (Figure 5 E, Supplementary Table 11). Only *TYR, TYRP1* and *DCT* were altered in response to acute zinc supplementation (Figure 5 D, Supplementary Table 7). TYR, TYRP1 and DCT are key to the production of melanin [21a] and pigmentary abnormalities show strong correlation with sub-RPE deposit formation and development of AMD [4a]. TYR catalyzes the production of melanin from tyrosine, in which L-DOPA is produced as an intermediate [96]. The function of TYRP1 is in the biosynthesis of melanin from tyrosine, whilst TYRP1 catalyzes the oxidation of 5–6-dihydroxyindole-2-carboxylic acid to an indole, whilst DCT catalyzes the conversion of L-dopachrome into 5-6-dihydroxyindole-2-carboxylic acid [96b]. These events lead to the activation of GPR143 signaling and may initiate several downstream effects, such PEDF, VEGF secretion and/or exosome release [21a]. Since we found an influence of zinc on the expression of the aforementioned genes, and given the data from literature above, zinc might also have an influence on GPR143 signaling. Surprisingly, acute zinc treatment resulted in down-regulation of the aforementioned genes, despite long-term zinc supplementation enhancing RPE pigmentation [8b]. At transcriptional level, long-term zinc supplementation significantly altered the expression of 18 out of the 21 pigmentation-related genes (Supplementary Table 11, [8d]), of which the majority were also down-regulated, except for *HPS5, HPS6, LYST*. These three upregulated genes are all related to intracellular trafficking, such as lysosomes and melanosomes [97].The negative effect of acute zinc supplementation on the gene expression of other pigmentation-related genes needs to be further investigated.

## 4. Conclusion

Primary hfRPE cultures that recapitulate the main phenotypes of aged RPE *in vivo* can help to dissect the molecular changes associated with RPE maturation and experimental manipulation, such as zinc supplementation. This cellular model provides an excellent platform for further preclinical studies to identify new treatment strategies for AMD. As reported *in vivo*, these cells retain a high degree of heterogeneity even after extended time in culture, which may help to understand the role of this heterogeneity in the human eyes. The identification of the transcriptional machinery, including transcriptional regulators ID1 and ID3 may help us to target pathways previously not considered for AMD. The data also shows that the differentiation of RPE into cells that resemble those *in vivo* requires an extended time in culture, and experimental manipulation will need to take this into account. The wide-ranging effects of zinc supplementation, from the regulation of housekeeping genes to very specific AMD associated transcripts builds confidence that this intervention could indeed be a suitable intervention strategy to slow the progression to advanced stage AMD, as suggested by the AREDS studies.

## 5. Methods

### 5.1. Retinal pigment epithelial (RPE) cell culture

Primary human fetal RPE cells (ScienCell, Carlsbad, USA) from one donor were purchased and used at passage number three (P3) for the complete study, in duplicates/triplicates with unknown clinical or genetic background. Cells were seeded onto Corning 6-well transwell inserts (10 μm thick polyester inserts with 0.4 μm pore size, 4*10^6^/cm^2^ pore density, Corning, Wiesbaden, Germany) in 125.000/cm2 in epithelial cell medium (EpiCM, ScienCell, Carlsbad, USA). After one week in culture, cell culture media was replaced with Miller medium with 1% FBS [98] and cells were cultured for two, nine, and nineteen weeks in duplicates. Two types of short-term zinc treatment were also conducted, where one-one extra replicates of untreated controls were taken for the two type of zinc treatment experimental setup. Either after one week or after eighteen weeks in culture, cell culture media was replaced with Miller medium with 1% FBS for an additional one week in the absence or presence of 125 μM externally added zinc (as zinc sulphate; Thermo Fisher Scientific, Waltham, USA) apically as well as basally, resulting in ~10 nM bio-available or free zinc [8b, 99]. The resulting replicates were the following: duplicates of zinc treated samples, triplicates of untreated controls at the two- and nineteen-week time-point and duplicates of untreated controls at nine-week time-point. Cellular differentiation was monitored through the development of cobblestone morphology and increase in pigmentation using light microscopy and the increase in trans-epithelial resistance (TEER) was measured by using the EVOM2 Epithelial Voltohmmeter and STX2 electrodes (World Precision Instruments, Sarasota, USA). At the sample collection time as above detailed, cells were washed with PBS (Thermo Fisher Scientific, Waltham, USA) two times for one minute, then cells were detached incubating cells with 0.15 % Trypsin-EDTA, thirty minutes, 37 °C. The trypsinization stopped using 100% FBS and trypsin neutralization solution (ScienCell, Carlsbad, USA). The obtained single cell suspensions were washed in PBS with 1% BSA (Thermo Fisher Scientific, Waltham, USA) 2 times, 5 minutes, 1000 rpm. After automatic cell counting (EVE, Thermo Fisher Scientific, Waltham, USA), 7 x 105 cells/ml were prepared, and the cells were kept on ice for a max ten minutes before proceeding with single cell RNA sequencing. In parallel to single cell sequencing, adjacent samples were fixed for fifteen min in 4% PFA (Merck, Darmstadt, Germany) diluted in PBS (Thermo Fisher Scientific, Waltham, USA) for immunofluorescence.

### 5.2. Experiment Overview

In this manuscript two separate scRNA-Seq experiments were performed from primary hfRPE cells from a single donor. In the initial scRNA-Seq run, samples originated from RPE cells cultured for two weeks (2W), nine weeks (9W) and nineteen weeks (19W) (Supplementary Figure 1A) in duplicates. Cells were collected from two wells at 2W, 9W and 19W. 7000 cells from each sample and were loaded on 10x Genomics Chromium v1.3 with target recovery of 4000. Libraries made from each sample were pooled and sequenced.

In the second run, samples originated from RPE cultures treated with a zinc-supplemented medium for one week either after: 1) one week in culture; or, 2) eighteen weeks in culture in duplicates. We also included one-one sample from untreated RPE culture in this run and the transcriptomic profiles were generated in a pooled fashion as described above. The actual cell recovery of both runs ranged from 3000 to 4000 in each well resulting in a total recovery of ~30,000 cells for the first run and ~15,000 for the second run. The raw scRNA-Seq data were processed using CellRanger v3.0.0. and then Seurat v3.1 to determine the heterogeneity of our specimens using unsupervised clustering, followed by annotation based on hierachical clustering of a pre-defined set of canonical RPE marker genes [11] (Supplementary Table 3). For further analysis, we initially analyzed our samples of untreated control RPE cultures from the two runs (triplicates for 2W and 19W and duplicates for 9W cultures) and then separately analyzed the duplicate samples of our zinc treated RPE cultures in comparison to the triplicate samples of untreated control RPE cultures of 2W and 19W).

### 5.3. scRNA-Seq

~ 7000 single cells per sample were processed with the Chromium system using the v3 single-cell reagent kit (10x Genomics, San Francisco, CA, USA). Barcoded libraries were pooled and sequenced on the NovaSeq platform (Illumina, San Diego, CA, USA), generating 150-base pair paired-end reads as per 10x Genomics recommendations, with >30000 reads per cell.

### 5.4. Bioinformatics

The raw scRNA-Seq data were processed using CellRanger version 3.0.0 (10X Genomics). The resulting filtered expression matrices were then imported into R for analyses using scRNA-Seq packages, Seurat (Version 3.1) (Stuart *et al*., 2019) and Monocle (Version 3.0) (Trapnell *et al*., 2014; Cao *et al*., 2019).

Cells were filtered to exclude those with <1000 or >8000 genes, or with >20% of counts aligned to mitochondrial genes, or >40% counts aligned to ribosomal genes. Cells passing QC were downsampled randomly to 1000 cells per sample to prevent over- or under-representation of any sample. Each sample was log-normalized using default Seurat parameters, with the top 3000 highly variable genes used for Seurat iterative pairwise integration. The integrated dataset was scaled to regress variance arising from read depth, and mitochondrial and ribosomal expression. Principal Component Analysis was then performed on the integrated dataset, and Seurat’s JackStraw function applied to determine the components used in UMAP and SNN clustering. Unsupervised clustering was run iteratively at resolutions ranging from 0.25 to 1, at increments of 0.25. At the highest resolution, a total of 13 clusters were detected. These clusters were observed in UMAP to form two overall, as-yet unannotated cell populations.

Using untreated cells only, the average expression for the clusters was determined for a set of 213 canonical RPE marker genes [11] (Supplementary Table 3), to which hierarchical clustering was applied. The clusters were segregated into two distinct branches, exhibiting characteristics of more- and less-differentiated RPE, which matched the distinction observed in UMAP. As such, the 13 unsupervised clusters were annotated to reflect these two overall cellular populations for downstream differential expression analysis.

Seurat’s Wilcoxon rank sum test was used for differential expression testing, using default FindMarkers parameters, with genes below 0.05 adjusted P value considered significantly differentially expressed.

Monocle 3 [100] was used for pseudotime analysis, for which downsampled count data was imported from Seurat, and independently processed and batch-corrected in Monocle using default parameters. A pseudotime trajectory graph was calculated and projected on the UMAP coordinates preserved from Seurat analysis for continuity. The data was filtered to focus on the main less-differentiated to more-differentiated pseudotemporal trajectory, by excluding small branches not contributing to the main trajectory. This was followed by graph-autocorrelation analysis to detect gene expression changes correlating with progress along the trajectory, filtered for significance at P-value and Q-value <0.05. Genes with expression significantly correlated with the trajectory were grouped into ‘modules’ of co-regulated genes, and the average expression of each gene module calculated across pseudotime.

### 5.5. Functional Classification Pathway and Network Analysis

For pathway and network analysis, we used the GeneAnalytics (https://ga.genecards.org/#input), and STRING (Search Tool for the Retrieval of Interacting Genes/Proteins) 11.0 (https://string-db.org/) [101] in combination with Cytoscape [102]. GeneAnalytics uses binomial distribution to test the null hypothesis that the queried genes are not over-represented within any super-path, GO term or compound in the GeneAnalytics data sources. The presented score in each section is a transformation of the resulting p-value, corrected for multiple comparisons using the false discovery rate (FDR) method; with higher scores indicating a better match. The bar color, indicating for the matching quality: high (dark green), medium (light green), low (beige) is common for all sections. STRING in combination with Cytoscape implements classification systems such as Gene Ontology, KEGG and systems based on high-throughput text-mining and the used reference dataset was the human genome. The identified functional protein association network was validated via text-mining, database information, co-expression, and experimental evidence.

### 5.6. Immunofluorescence

For immunofluorescence analysis, the cells on the transwell membrane were permeabilized in 0.5% Triton-X (Merck, Darmstadt, Germany) in PBS for ten minutes at 4 °C and then washed in 0.1% Tween20 in PBS (PBST) (Merck, Darmstadt, Germany) and blocked with 5% goat sera (Merck, Darmstadt, Germany) in PBST with for one hour at room temperature. Samples were then incubated with primary antibodies for overnight: *COL1A1* (Abcam plc, Cambridge, UK, dilution 1:200) and *RPE65* (Merck Millipore, Darmstadt, Germany, 1:50), diluted in PBST containing 1% goat sera. Following washing with PBST, the samples were incubated with secondary antibodies in 1:200 in PBST with 1% goat sera for one hour in the dark at room temperature. Samples were washed with PBST for five minutes, then with PBS. Cell nuclei were then labelled with DAPI (Thermo Fisher Scientific, Waltham, USA) diluted 1:1,000 in PBS. Finally, samples were mounted onto Menzel-Glaser slides (Thermo Fisher Scientific, Waltham, USA) in Vectashield (Vector Laboratories, Burlingame, USA). For negative control, the primary antibody labelling was omitted. Cells were visualized using a Leica SP8 confocal microscope (Leica, Wetzlar, Germany). Images were obtained and analysed with Leica Application Suite X Image software (Leica, Wetzlar, Germany).

## Acknowledgements

The authors gratefully acknowledge the QUB Genomics Core Facility for their expertise and assistance in this work.

## Data availability statement

The data that support the findings of this study will be openly available in the GEO database public repository upon publication, that does not issue DOIs.

## Author Contributions

Conceptualization, E.E., O.C., E.K., D.S. and I.L.; Methodology, E.E. and O.C, Investigation, E.E., O.C., D.S. and I.L.; Formal Analysis and Visualization, E.E., O.C., C.K., J.P.S.G., D.S. and I.L.; Writing—Original Draft, E.E. and I.L.; Writing—Review and Editing, E.E., O.C., E.K., B. McK., J.P.SG., A.A.B., D.S. and I.L.; Supervision and Funding Acquisition I.L., D.S. and A.A.B; Project administration I.L. All authors have read and agreed to the published version of the manuscript.

## Supplementary Figures

**Supplementary Figure 1.:**
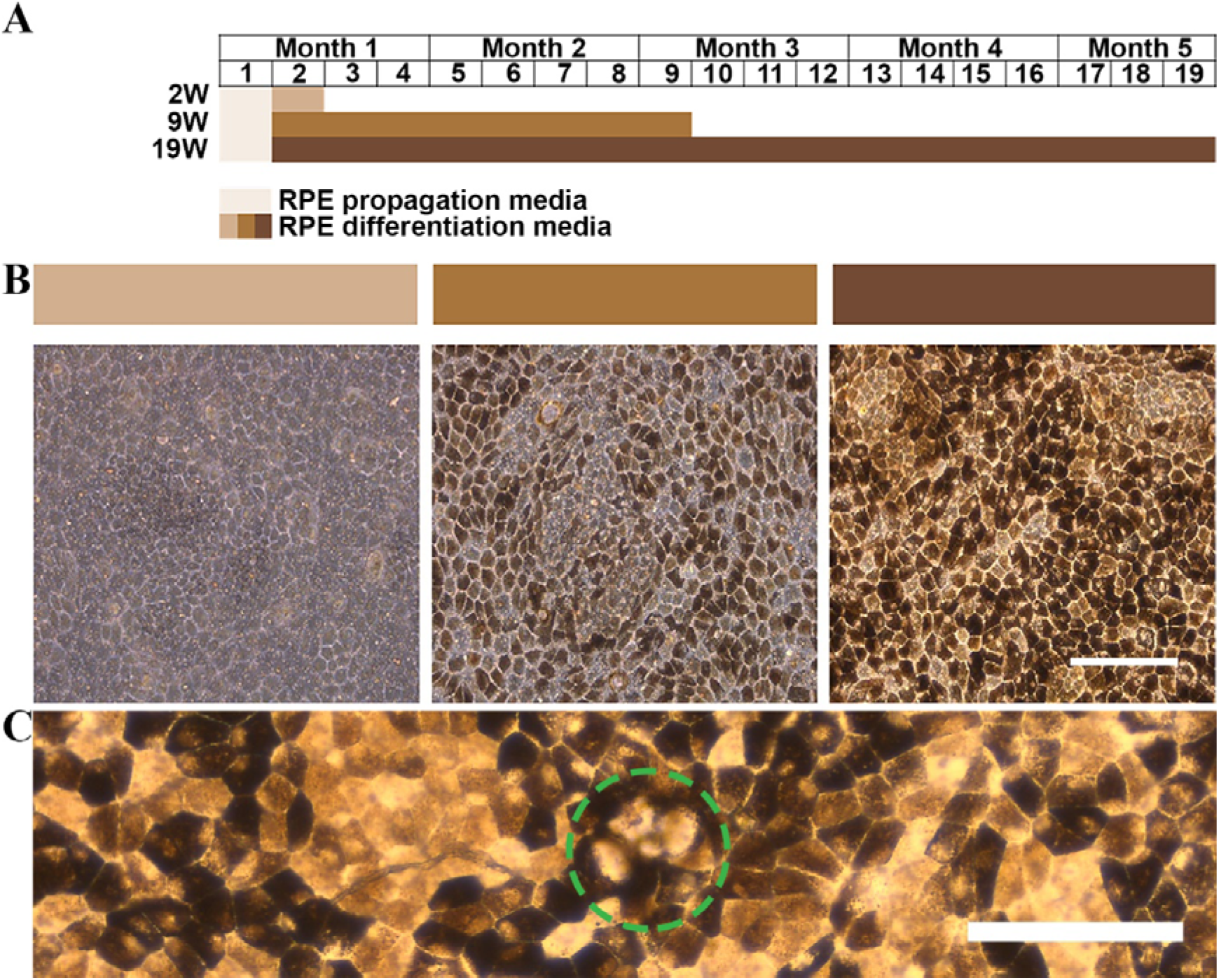
*In vitro* primary fetal RPE culture. Experimental design of followup maturing RPE *in vitro* (A), pigmentary changes of *in vitro* RPE over time (B) and representative deposit accumulation in 19 weeks old cells (C).

**Supplementary Figure 2.:**
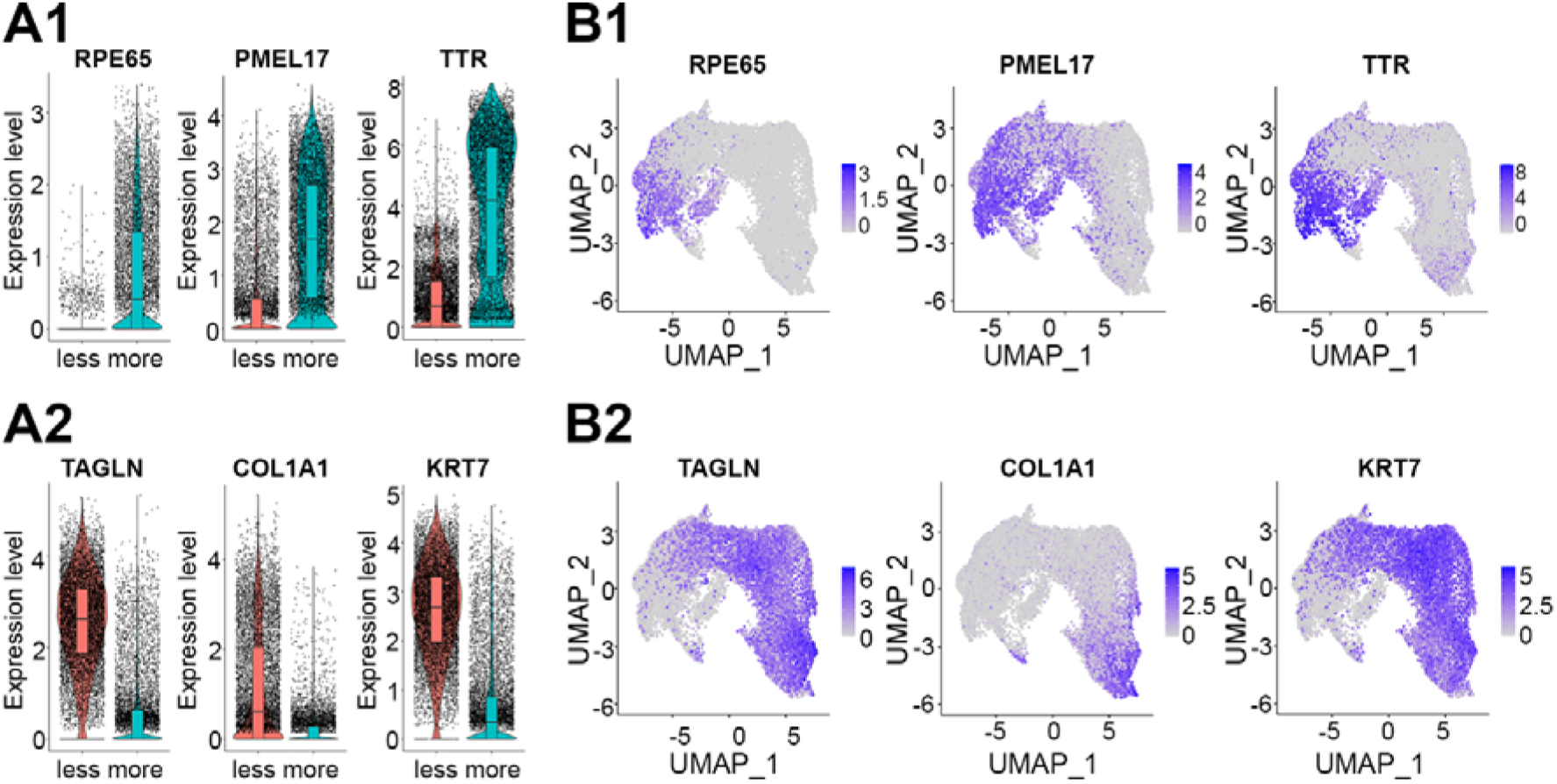
Comparing gene expression patterns of more and less differentiated groups. Three highly expressed marker genes RPE65, PMEL and TTR showed higher expression in the more differentiated cells (A1). Expression of TAGLN, COL1A1and KRT7 were higher in less differentiated cells (A2). The distribution of cells expressing these genes on the UMAP shows the enrichment of cells in the more (B1) or less (B2) differentiated groups. Note: The genes individually did not place every cell into the appropriate groups.

**Supplementary Figure 3.:**
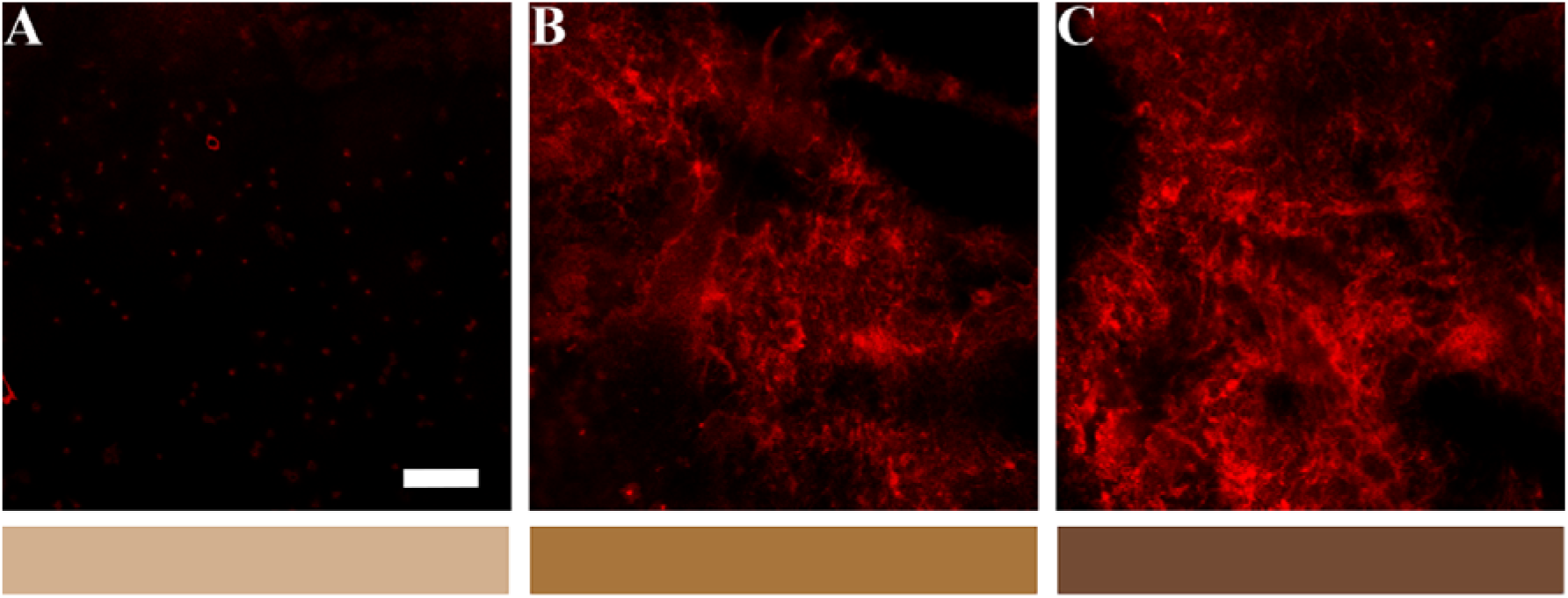
Basal immunolabeling of COL1A1 protein increased in the RPE culture experiments. A, short-term, B, medium-term, C, long-term culture. Scalebar is 10 um.

**Supplementary Figure 4.:**
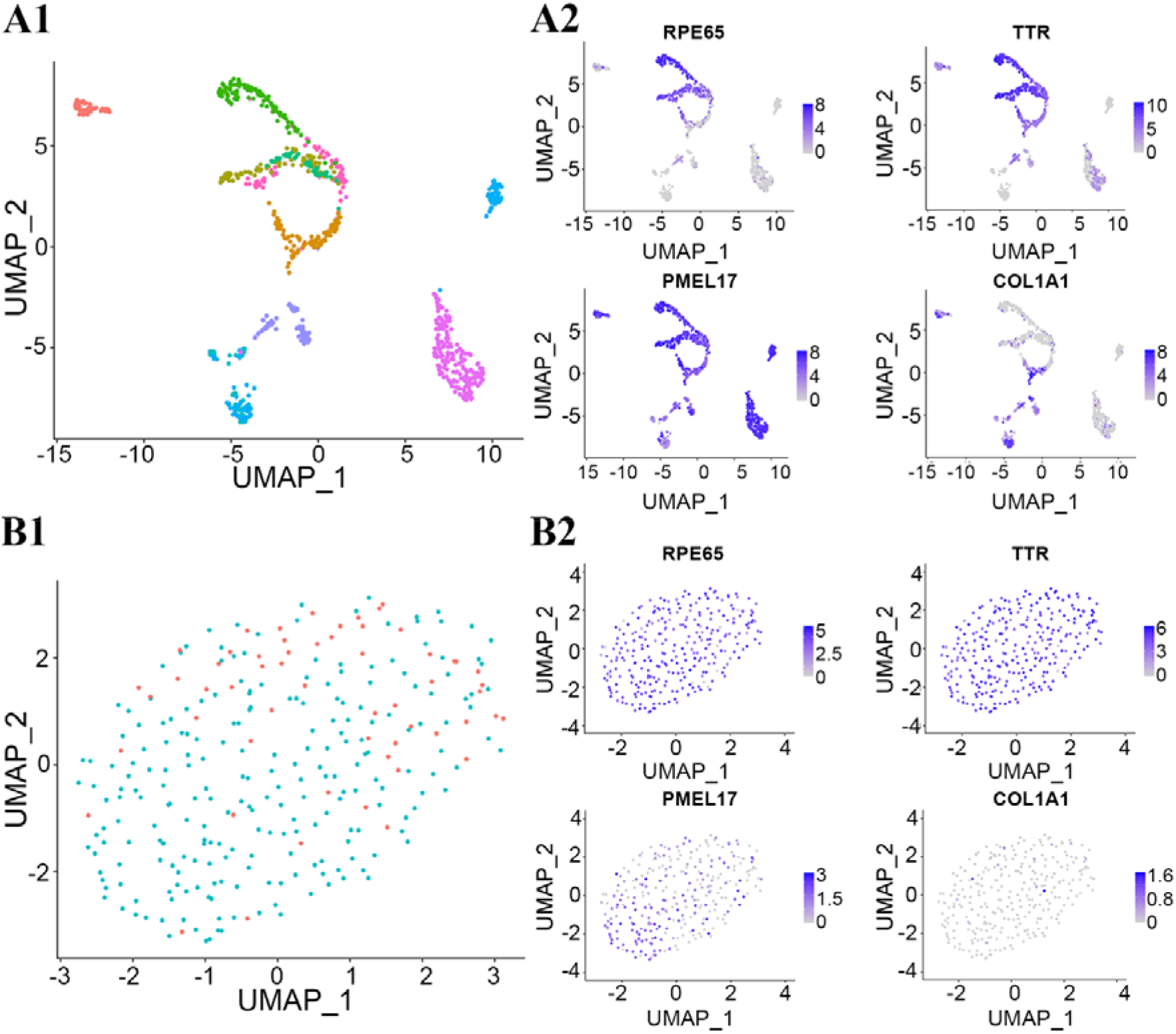
Gene expressional patterns of single-cell populations in the developmental and adult RPE *ex vivo*. Heterogeneity of RPE at single-cell level on *ex vivo* embryonic developmental RPE (A1) and of adult RPE (B1) and gene expression pattern of RPE65, TTR, PMEL and COL1A1 during development (A2) and in adult stage (B2) of RPE.

